# Kinetic Modeling of Covalent Inhibition: Effects of Rapidly Fluctuating Intermediate States

**DOI:** 10.1101/2025.05.28.656658

**Authors:** Kyle Ghaby, Benoît Roux

## Abstract

There is increasing interest in the discovery of small-molecule inhibitors that form covalent bonds with their targets for therapeutic applications. Nevertheless, identifying clear rational design principles remains challenging because the action of these molecules cannot be understood as common noncovalent inhibitors. Conventional kinetic models often reduce the binding of covalent inhibitors to a two-step irreversible process, overlooking rapid complex dynamics of the associated unlinked inhibitor before the formation of the covalent bond with its target. In the present analysis, we expand the intermediate state into two conformations—reactive (E·I) and nonreactive (E··I). To illustrate the consequences of such simplification, the expanded kinetic model can be reduced to an effective two-step scheme expressed in terms of the equilibrium probability of the unlinked inhibitor to form either conformation. A mass-action-based numerical workflow is implemented to simulate time-dependent kinetics, overcoming the common limitations of empirical models. The numerical workflow helps relate microscopic states observed in molecular dynamics simulations to macroscopic observables like EC_50_ and the apparent rate of covalent inhibition, showing the impact of transient intermediates on dissociation rates and potency. The proposed framework refines the interpretation of dose-response data, aiding medicinal chemists in optimizing covalent inhibitors and provides a quantitative platform for relating molecular conformational distributions to empirical parameters.

**TOC Graphic:** 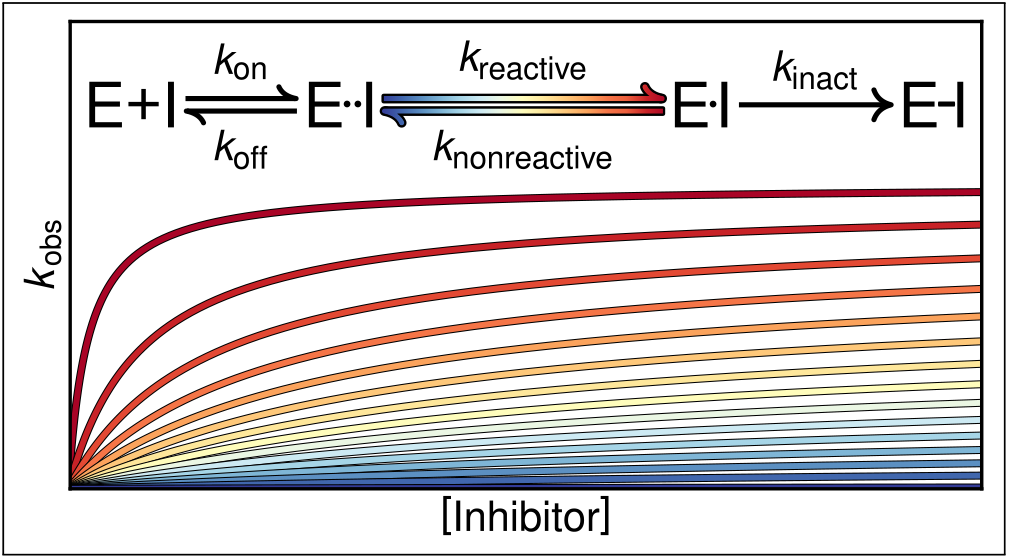

## Introduction

For a long time, inhibitory small-molecules that form covalent bonds with their targets, *i*.*e*. covalent inhibitors, were largely eschewed in drug development due to concerns over off-target effects, assay and delivery interference, and metabolitic toxicity.^1–8^ With the improvements in drug design and optimization over the past couple decades, targeted covalent inhibition is becoming a promising and widely recognized strategy for exceeding the potency and selectivity of noncovalent inhibition.^9–26^ These enhancements should lead to improvements in the translation of *in vitro* to *in vivo* efficacy and safety profile.^27,28^ Covalent inhibitors are also useful as probes for chemoproteomic applications.^29^

Nevertheless, the rational design of covalent inhibitors remains extremely challenging. Today’s dominant strategy to design a covalent inhibitor is to start from a molecular scaffold that displays some reversible binding affinity to a protein of interest and attach to it a reactive functional group or “warhead” that targets a specific residue in the binding site.^24,30–38^ Conventional docking methods developed to deal with noncovalent protein–ligand interactions have been adapted to model the binding of a covalent ligand with the protein.^39–45^ It is commonly assumed that poses with the warhead in close proximity to the target residue can be used to predict a covalently-bound adduct, and computational workflows based on these ideas have been proposed.^34–36,46–50^ Nevertheless, there is a wide range of issues. In particular, conventional docking often falls short when a protein target is flexible and can adopt multiple distinct conformations. It is also worth emphasizing that the mere presence of a reactive residue near a ligand binding site is not sufficient to guarantee a successful covalent strategy; the local chemical environment of the target residue must be carefully considered to determine the suitability of this approach.^51^ Dynamics before the covalent bond is formed is also thought to play a key role.^52,53^ Covalent inhibition ultimately depends on a chemical reaction taking place in the confined environment of a protein binding pocket. Residues near the warhead, in turn, may act as catalysts or steric obstacles at various steps along the reaction pathway. The situation is very different than for a reaction occurring in bulk solution. Despite their short lifetime, the rapidly fluctuating intermediate conformational states at play in all of these issues have a profound and complex impact on the apparent reaction rate.

The purpose of kinetic models, in the context of a drug development project, is to help draw accurate lessons from experimental observations in order to make judicious decisions to orient the design strategy in a productive direction. Due to the limited resolution, kinetic measurements of covalent inhibition have commonly been interpreted in terms a simple two-step mechanism.^54–56^ The simple picture is that the inhibitor initially diffuses in the bulk solution to encounter the kinase enzyme and associate with the active site, followed by the irreversible inactivation of the enzyme due to the formation of the covalent adduct. A critical issue with such a minimal two-step model is that it may obscure important microscopic factors that are at play, which in turn may misdirect the design strategy. For example, kinetic data interpreted in light of the two-step model tend to ascribe the apparent rate of inactivation to the final chemical step that leads to the formation of the covalent linkage. Thus, while a slow inactivation rate constant may be caused by a low probability of a reaction-ready pose by the unlinked ligand occupying the binding site, the observation of an apparent slow rate might erroneously be attributed to the poor reactivity of the warhead, thereby leading the design strategy in a wrong direction.

While the mathematical simplicity of a model is an important advantage to analyze experimental data, it should not be a determining factor. Computational methods for numerically predicting molecular behavior and solving complex kinetic schemes have grown much more accessible and reliable. The precision of analysis depends on the precision of the kinetic scheme. Also of interest, the fluctuations predicted by classical molecular dynamics (MD) simulations based on atomic models can provide quantitative information that can be incorporated into a kinetic scheme to affect the observable covalent occupancy. A recent computational study of the irreversible covalent inhibitor of Bruton’s tyrosine kinase (BTK) ibrutinib illustrates these ideas.^52^ Classical MD simulations of the unlinked compound associated with the kinase binding pocket showed that only a small fraction (less than 3%) of the configurations were reaction-ready despite the fact that the inhibitor remains very close to the crystallographic binding pose. Quantitative agreement with experimental data was achieved only once this information was combined with the result of hybrid QM/MM calculation of the covalent reaction. The kinetic effects of rapidly fluctuating intermediate conformational states have been characterized and discussed in other fields,^57–59^ but has not been extensively considered in covalent inhibition.

Owing to their growing significance, it is important to develop more accurate and more realistic mathematical models of covalent inhibitors.^60–68^ It is the central goal of the present work to help clarify the impact of rapid conformational fluctuations on the apparent potency of covalent inhibitors, which should hopefully lead to more effective design strategies.

## Theoretical Framework

### Kinetic schemes

At the simplest level, the kinetics of irreversible covalent inhibition are often described as a two-step mechanism shown in Scheme 1,^54–56^

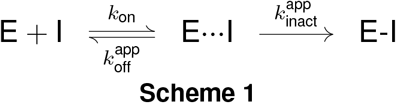

where *k*_on_ is the bimolecular association rate, 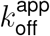 is the apparent dissociate rate, and 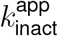 is the apparent inactivation rate constant. Initially, the inhibitor (I) diffuses in the bulk solution to encounter the kinase enzyme (E) and associate with the active site, resulting in the reversible noncovalent complex (E···I). The final step describes the inactivation of the enzyme as an irreversible process that results in the covalent adduct (E-I). Intermediates such as noncovalent fluctuations and the quantum-mechanical mechanisms of bond formation are combined in E···I and accounted for in 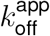 and 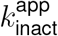. We explicitly attach the label ‘app” to these rates because they implicitly subsume additional processes, as will be clarified in the following.

Scheme 2 splits the state E···I into nonreactive (E··I) and reactive (E·I) intermediate states, where the reactive intermediate state meets some distance and orientation requirements necessary for the progress of the covalent reaction between the warhead and the target residue. For the sake of simplicity, it is assumed that only the nonreactive state can dissociate.

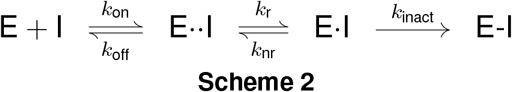

The equilibrium probability of the unlinked inhibitor to form the either reactive (*p*_r_) or nonreactive complex (*p*_nr_) can be defined from the species concentrations as,

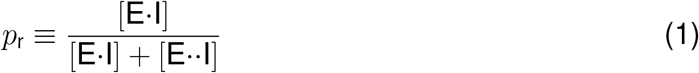

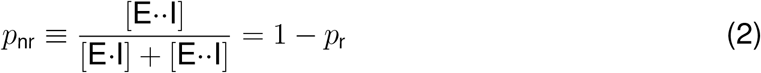

Classical MD simulations indicate that the fluctuations of the unlinked inhibitor between the reactive and nonreactive pose may take place on a timescale of microseconds.^52^ This is significantly faster than the other transitions, whose rate constants are typically on the order of minutes or even hours. Since these states are rapidly equilibrated relative to the rest of the scheme,

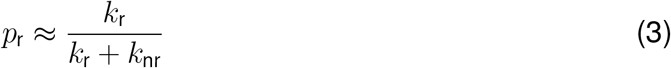

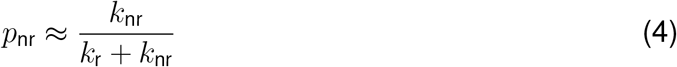

and Scheme 2 can be reduced and made equivalent to a two-step system, as shown in Scheme 3,

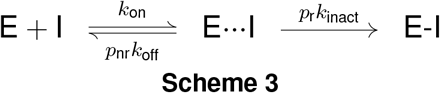

where the apparent dissociation rate constant 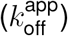 equals *p*_nr_*k*_off_, and the apparent inactivation rate constant 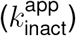 equals *p*_r_*k*_inact_. This equivalence was verified with numerical simulations, *i*.*e*., for the cases examined in the present study, the results obtained by solving the ODEs numerically for Scheme 2 or from Scheme 3 were identical. To highlight the effects of *p*_r_ in the absence of the irreversible covalent step from Scheme 2, it is also of interest to consider a non-covalent version as Scheme 4,

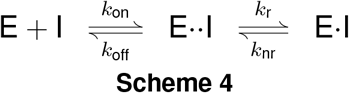

which is also useful for illustrating the influence of rapid intermediates on the residence time (*τ*) of a bound ligand.

### Simulation methodology

The time course of species concentrations in a well-defined (*i*.*e*. the rate constants and stoichiometry are known or parametric for each transition), Markovian kinetic scheme can be solved deterministically by integrating a set of ordinary differential equations (ODEs) for concentration with respect to time.^69^ These ODEs are derived from the law of mass action and formulated based on the stoichiometric coefficients and rate constants of each transition in the scheme. This fast, flexible, and straightforward framework adequately models the *in vitro* kinetics^70^ of covalent inhibition.

Let *C*_*S*_(*t*) represent the concentration of species *S* at time *t*. The time derivative *dC*_*S*_*/dt* is given by the sum of contributions from all transitions involving species *S*,

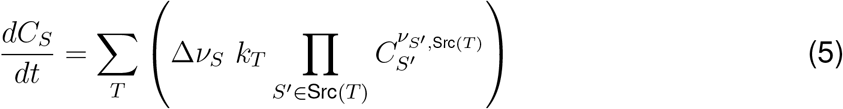

where *k*_*T*_ is the rate constant of transition *T*, and Δ*ν*_*S*_ is the difference between the stoichiometric coefficients of species *S* as a target and as a source in transition *T* (*i*.*e. ν*_*S*,Tgt(*T*)_ − *ν*_*S*,Src(*T*)_).

A system of these rate laws with any degree of (non)linearity can be expressed and solved in matrix notation as,

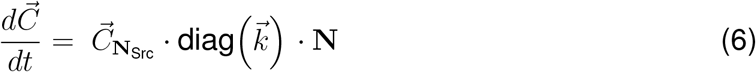

The complete stoichiometry matrix, N, is the difference between the target and source stoichiometry matrices (*i*.*e*. N = N_Tgt_ ™ N_Src_). The stoichiometric matrices have *m* rows corresponding to the transitions and *n* columns corresponding to the species. 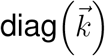 is the diagonal matrix constructed from the vector of transition rate constants. The adjusted concentration vector, 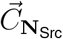, is the column-wise (*i*.*e*. species-wise) product of the element-wise exponentiation of the concentration vector with the source stoichiometry matrix, *i*.*e*.,

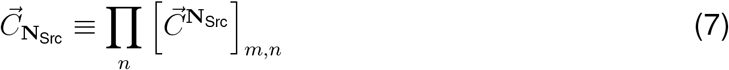

eq 6 is numerically integrated over time with the scipy implementation of LSODA.^71–73^ This vectorized rate function together with the rest of the infrastructure for modeling the schemes comprises GeKiM, an in-house and open-source Python package.

A Jupyter notebook to reproduce the simulations and plots herein is available on Github. The numerical values of the rate constants used in the simulations were chosen to reflect the regime commonly observed for covalent inhibitors, where *k*_inact_ ≪ *k*_off_ *< k*_on_[I]_0_.^74^

## Results

### Noncovalent vs covalent inhibition

Time courses of the total occupied populations of Scheme 2 and its noncovalent version Scheme 4 are compared in Figure 1 for various probabilities of a reactive conformation, *p*_r_. The purpose of Scheme 4 is to highlight the effects of *p*_r_ in the absence of the irreversible covalent step from Scheme 2. *p*_r_ has a similar impact on the occupancy of both of the schemes. The noncovalent time courses demonstrate how *p*_r_ can compensate for poor affinity and limit maximal occupancy. The irreversible covalent time courses demonstrate the dependence of covalent action rate on *p*_r_. Reversible covalent inhibition falls somewhere in between, depending on the ratio of covalent forward and backward rate constants.

**Figure 1:**
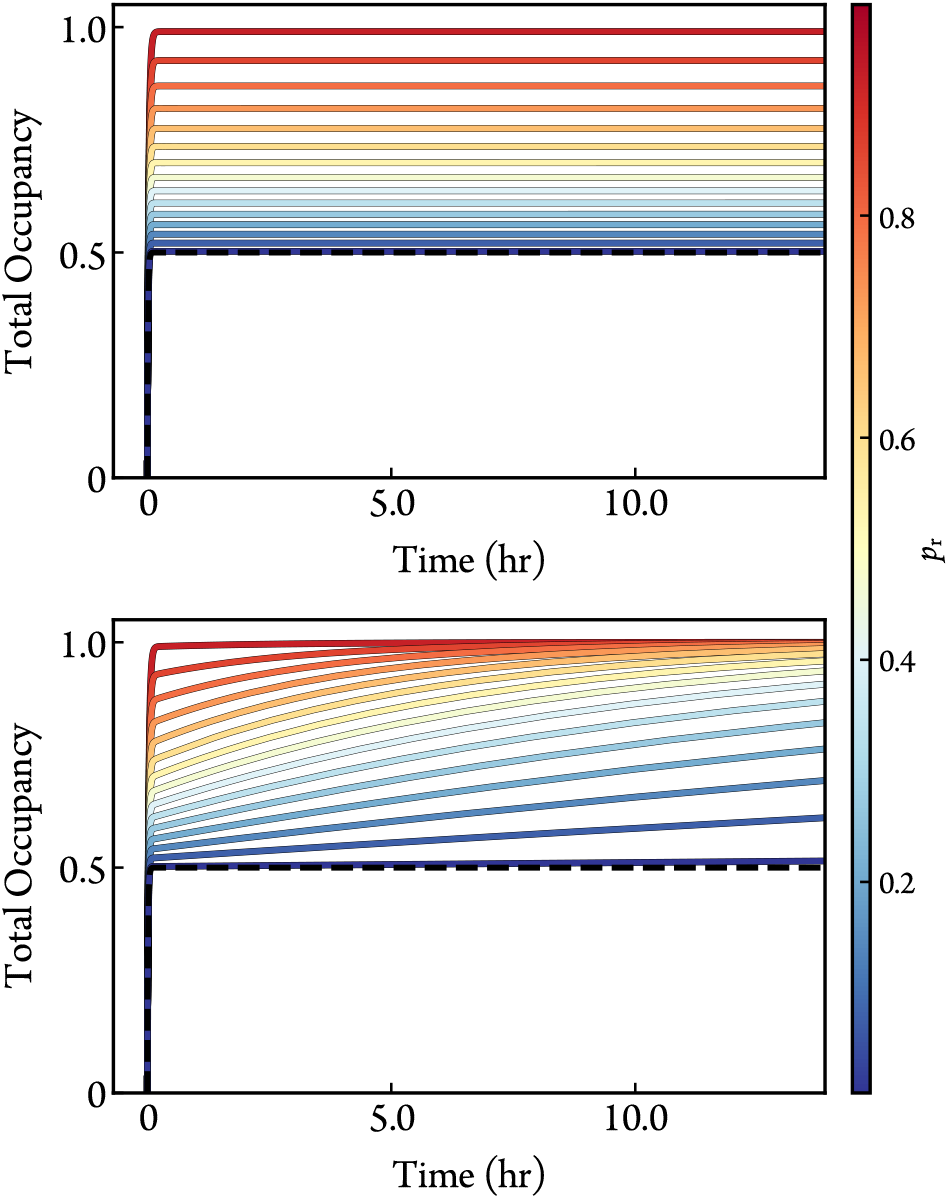
Simulated time courses of the total occupancy from Scheme 2 (bottom) and its noncovalent version from Scheme 4 (top) for different values of *p*_r_ with [E]_0_ = 1 nM, [I]_0_ = 1 µM, *k*_on_ = 1× 10^−2^ µM^−1^s^−1^, *k*_off_ = 1 ×10^−2^ s^−1^, and *k*_inact_ = 1 10^−4^ s^−1^. The occupancy of a one-step scheme is also included (black-dashed) to illustrate the contribution of binding affinity, *k*_off_*/k*_on_.

As shown in Figure 2, the propensity to adopt a conformation in which the ligand cannot dissociate significantly increases the lifetime of binding despite the short lifespan of the conformations themselves. The residence time derived from Scheme 4 can be expressed using the rapid-equilibrium approximation as,

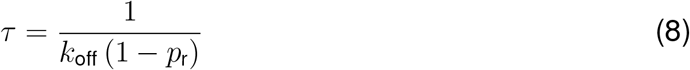

**Figure 2:**
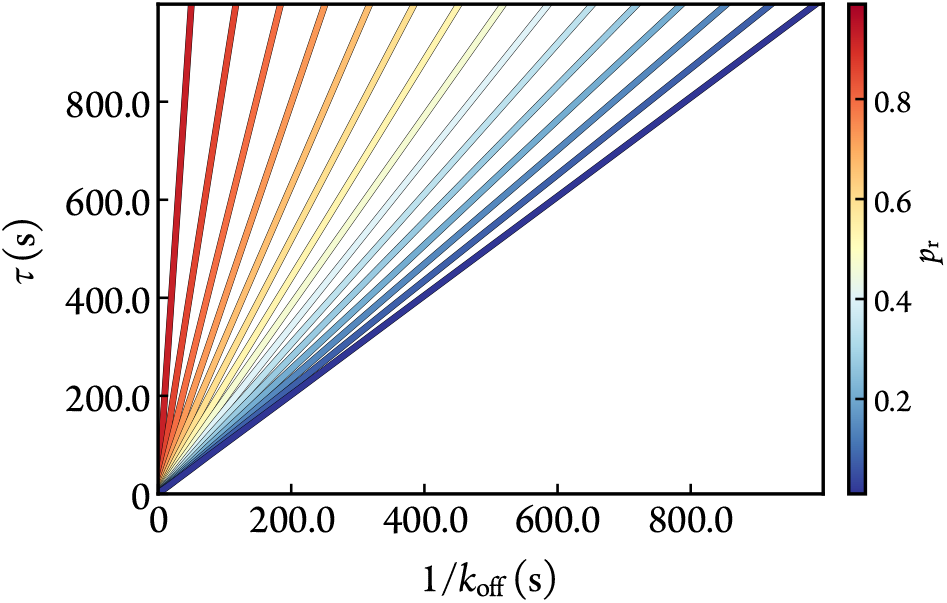
Noncovalent residence time (*τ*) of a bound ligand as a function of *k*_off_ and *p*_r_. [E]_0_ = 1 nM, [I]_0_ = 1 µM, and *k*_on_ = 1 ×10^−2^ µM^−1^s^−1^. The residence times calculated analytically from eq 8 or numerically from simulations of Scheme 4 are the same.

### Effects on observable outcomes

#### Dose-rate analysis

The kinetics of covalent inhibitors are often studied in a pseudo-first order regime in the spirit of Michaelis-Menten kinetics,^75,76^ with an excess of either inhibitor or enzyme. For the sake of convention, we will treat the inhibitor as the saturating species, so the biomolecular step can be linearly approximated as shown in Scheme 5.

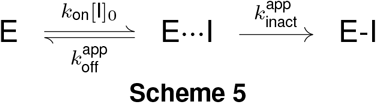

In conventional experimental procedures of scoring an irreversible inhibitor’s on-target behavior,^56^ the observed rate constant of covalent bond formation, *k*_obs_, is first obtained by fitting the fractional population of the covalently-bound state (*i*.*e*. covalent occupancy—CO) to the pseudo-first order growth factor,

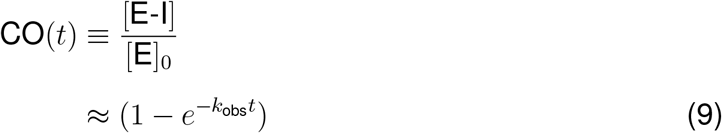

at various inhibitor concentrations. The variation of *k*_obs_ as a function of initial inhibitor concentration, [I]_0_, is then fitted to the Michaelis-Menten analog,

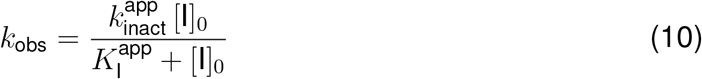

The apparent rate constant of inactivation, 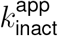, is the maximal observed rate constant that occurs when the entire free enzyme population rapidly transitions into the reversible complex state. This limit is approached when 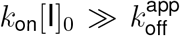. The apparent inactivation constant, 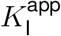, is the concentration of the inhibitor that results in a half-maximal observed rate constant, and it can be expressed as,

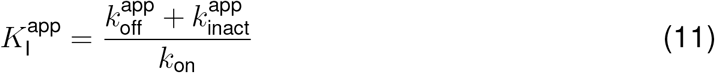

A reaction may appear one-step in both non-saturating ([I]_*0*_ *< K*_I_) and saturating ([I]_0_ ≫ *K*_I_) conditions. These one-step approximations are distinguishable by the *y*-intercept, where an intercept of 0 corresponds to a slope of *k*_obs_ over [I]_*0*_ equivalent to the inactivation efficiency (*i*.*e*. the potency of the irreversible inhibitor).^77^ For the three-step paradigm (Scheme 2),

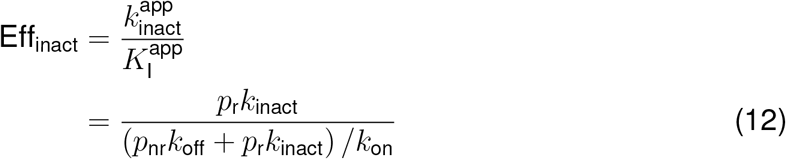

The probability distribution of the intermediates has a significant role in the potency of inhibition despite the non-limiting magnitude of the arming and disarming rate constants. From an empirical perspective, the probability distribution between conformations limits inhibition by linearly scaling the maximal observed rate constant. 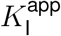 also depends linearly on *p*_r_, with,

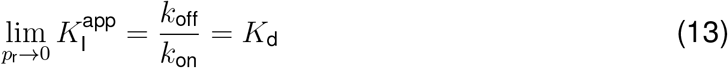

and

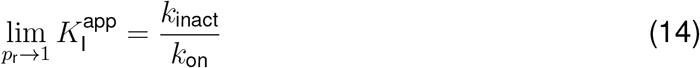

Rearranging eq 14 for *k*_on_ shows that Eff_inact_ approaches *k*_on_ as *p*_r_ approaches 1; a reminder that simplifying eq 12 at 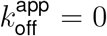 yields *k*_on_. The dramatic relationship between simulated empirical data from the numerical solution to the ODEs of Scheme 3 and *p*_r_ is illustrated in Figure 3.

**Figure 3:**
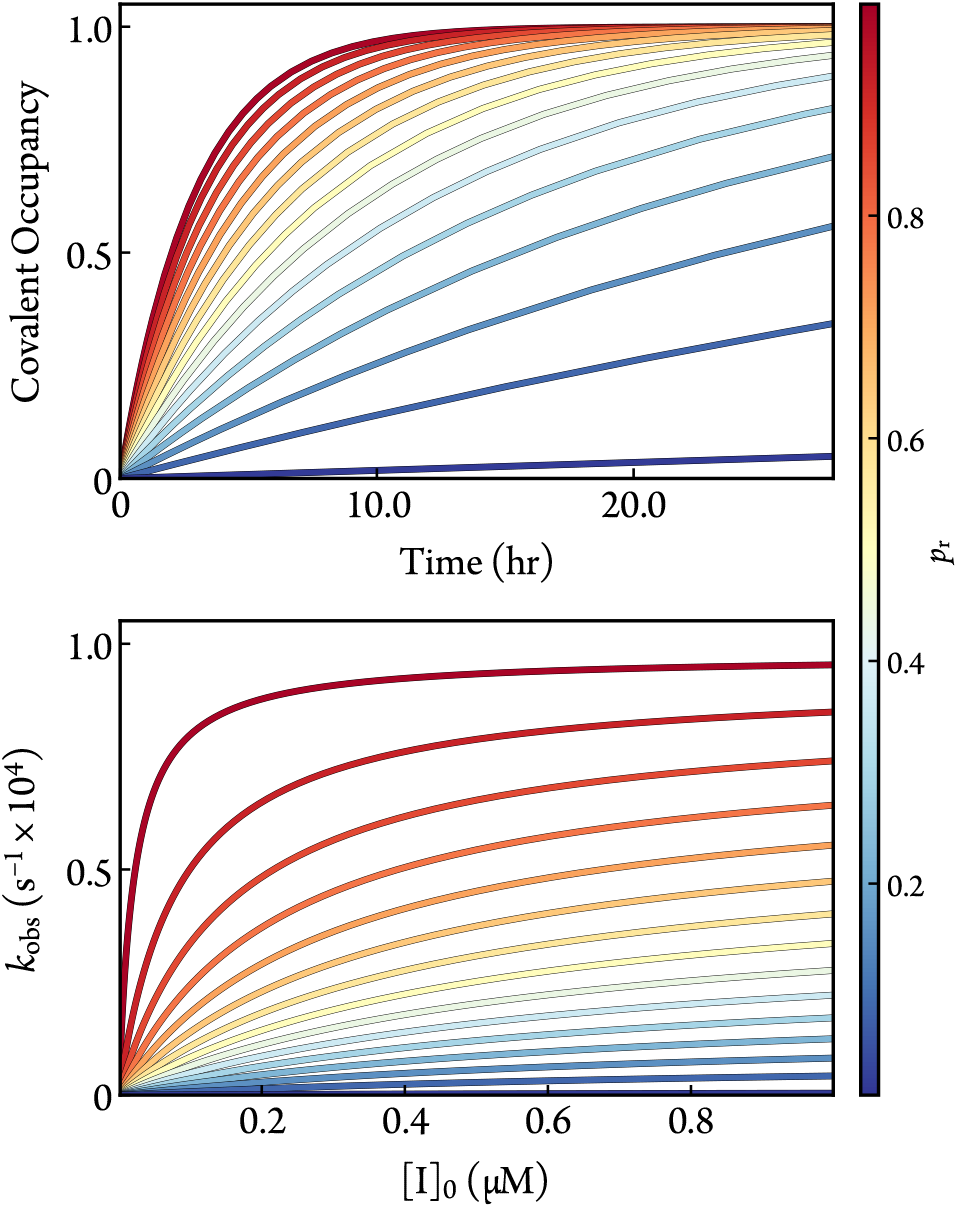
Simulated time-responses of covalent occupancy ([I]_0_ = 1 µM) (top) and dose-rate (bottom) data for different values of *p*_r_ with [E]_0_ = 1 nM, *k*_on_ = 1× 10^−2^ µM^−1^s^−1^, *k*_off_ = 1× 10^−2^ s^−1^, and *k*_inact_ = 1× 10^−4^ s^−1^. The ODEs of Scheme 3 were solved numerically for the time-responses (top), and the fraction of the covalent complex was fit for *k*_obs_ with eq 9 (bottom).

### Dose-response analysis

The dose-dependence of irreversible kinetics can alternatively be examined with a dose-response curve. Unlike reversible kinetics, the dose-response must be examined under non-equilibrium conditions where the reaction is stopped before completion. For simplicity, we will only consider noncompetitive experimental procedures where some proportional marker of covalent occupancy, *i*.*e*. response, would be measured at the same timepoint for an array of saturating ligand concentrations. While the numerical simulations of the ODEs provides the most faithful representation of a given kinetic scheme, the familiar Hill equation,^78^

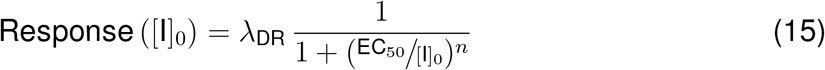

can serve as a useful shorthand analytical framework for describing the dose-dependence data because it produces a simple sigmoidal curve with clearly identified parameters. In eq 15, *λ*_DR_ represents a scaling factor to match the arbitrary amplitude of the data, EC_*50*_ represents the dose that yields half the maximal scaled response, and *n* represents the Hill coefficient.

The Hill equation works exceedingly well for reversible one-step systems measured at equilibrium, where the EC_*50*_ approximately equals the apparent binding affinity, 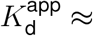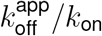 (*i*.*e*. the apparent inhibition constant, 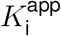). However, special attention is required in the case of a covalent inhibitor because the covalent occupancy becomes truly stable only when the entire enzyme population has been irreversibly occupied. Thus, the value of EC_*50*_ is time-dependent and its kinetic significance is ambiguous.^61,69^ In the following, we assess those kinetic parameters in terms of 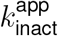 and 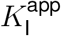 for the purpose of describing dose-dependence data for a covalent inhibitor.

First, the expression must be properly normalized to covalent occupancy. This can be done with the exponential growth factor for approximating a time-response curve of covalent occupancy (eq 9). Recall the empirical definition of 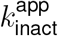 : the maximal observed rate constant that occurs when *k*_on_[I] is large enough to instantly and steadily force the entire (remaining) protein population into the reversibly-bound state. The Hill equation is then scaled by the maximal covalent occupancy achieved within the dose-response time limit (*t*_DR_) to get,

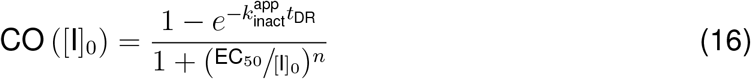

Thus, the amplitude of eq 16, *i*.*e*. the limit of covalent occupancy given a set time and an infinitely high dose, is solely dependent on 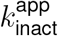 (or by extension *k*_inact_ and *p*_r_).

To characterize the impact of the kinetic parameters on the location of the inflection point of eq 16, *i*.*e*. the dose required for half-maximal covalent occupancy within the time limit, we solve for the isosurface of EC_50_ in terms of 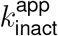 and 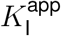. We start by solving for *k*_obs_ from the concentration at half maximum inhibition, defining [I]_0_ = EC_50_. Using the definition of the EC_50_, a relationship of familiar first-order growth factors can be expressed at this condition,

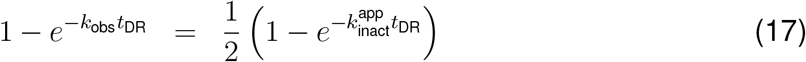

We then solve for *k*_obs_,

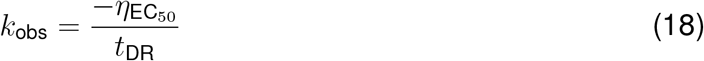

where

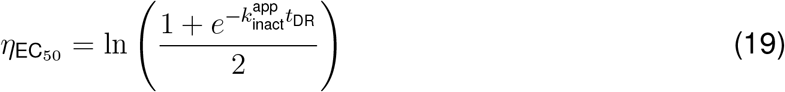

Then, we substitute *k*_obs_ with the Michaelis-Menten analog (eq 10) when [I]_0_ = EC_50_ to get the approximation,

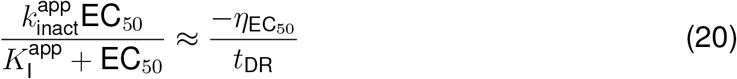

Finally, we solve for the EC_50_,

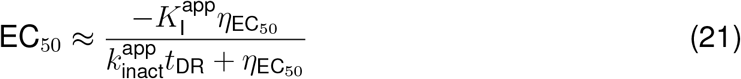

As is expected from reversible one-step kinetics, the EC_50_ approaches the reversible binding affinity, *K*_d_, as 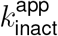 approaches 0. This expression of the EC_50_ is substituted into eq 16 to guide the understanding of dose-dependence with respect to the microscopic kinetic parameters of irreversible binding.

Still, this expression is an approximation of the kinetics. Covalent occupancy calculated from eq 16 and ODE-based simulations of Scheme 3 is compared in Figure 4 for three demonstrative cases. The same rate constants are used in both solutions for each case, and the value of *n* in eq 16 is fit to the simulated covalent occupancy. eq 16 deviates from the numerical covalent occupancy in Cases 1 and 3. Case 1 represents the regime where 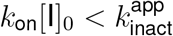. In this regime, a single dose-response is likely not enough to resolve these rate constants anyway. Case 3 includes nonlinear effects from the bimolecular association step. The nonlinearity violates the first-order approximation used to derive eq 21, and therefore leads to a poor analytical prediction of the EC_50_.

**Figure 4:**
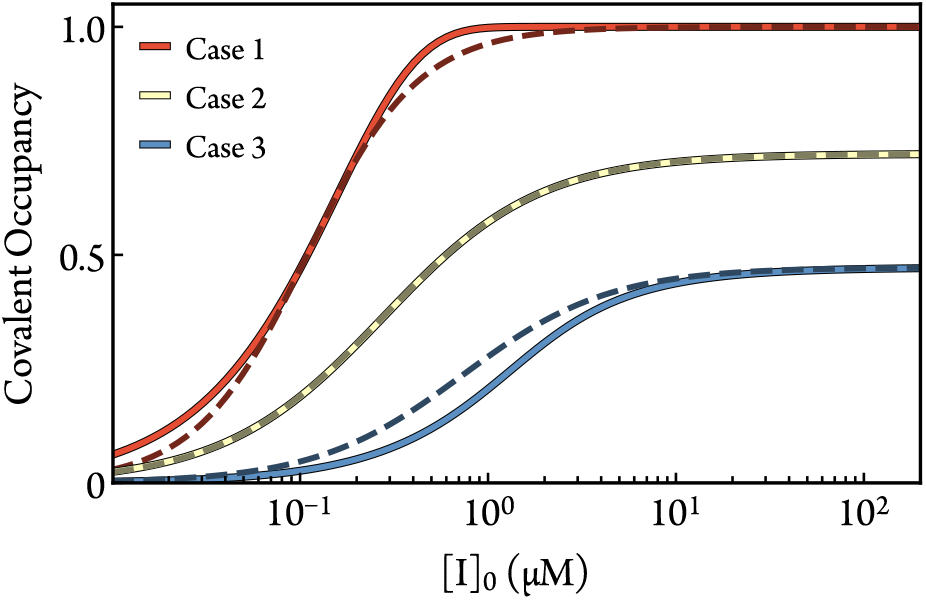
Comparisons between eq 16 (darkened dashed line) and ODE-based numerical simulations of Scheme 3 (solid line) for three representative cases. In Case 2, the parameters are *t*_DR_ = 7.1 h, [E]_0_ = 1 nM, *k*_on_ = 1 × 10^−2^ µM^−1^s^−1^, 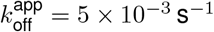, and 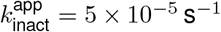. The altered parameters for Case 1 are *k*_on_ = 5 × 10^−4^ µM^-1^ s^−1^ and 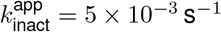. The altered parameters for Case 3 are [E]_0_ = 1 µM, *k*_on_ = 5 × 10^−4^ µM^−1^s^−1^, and 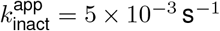. The fitted values of *n* for cases 1, 2, and 3 are 1.47, 1.03, and 1.11, respectively.

According to the present analysis, the EC_50_ depends both on 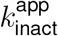 and 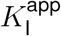. A smaller EC_50_ can either be attributed to a smaller *K*_d_ (or faster *k*_on_ for systems lacking reversible equilibrium), larger *p*_r_, or faster *k*_inact_. Therefore, the EC_50_ by itself is a vague and possibly misleading parameter for evaluating irreversible systems. This mathematical analysis is illustrated with numerical kinetic simulations in Figure 5.

**Figure 5:**
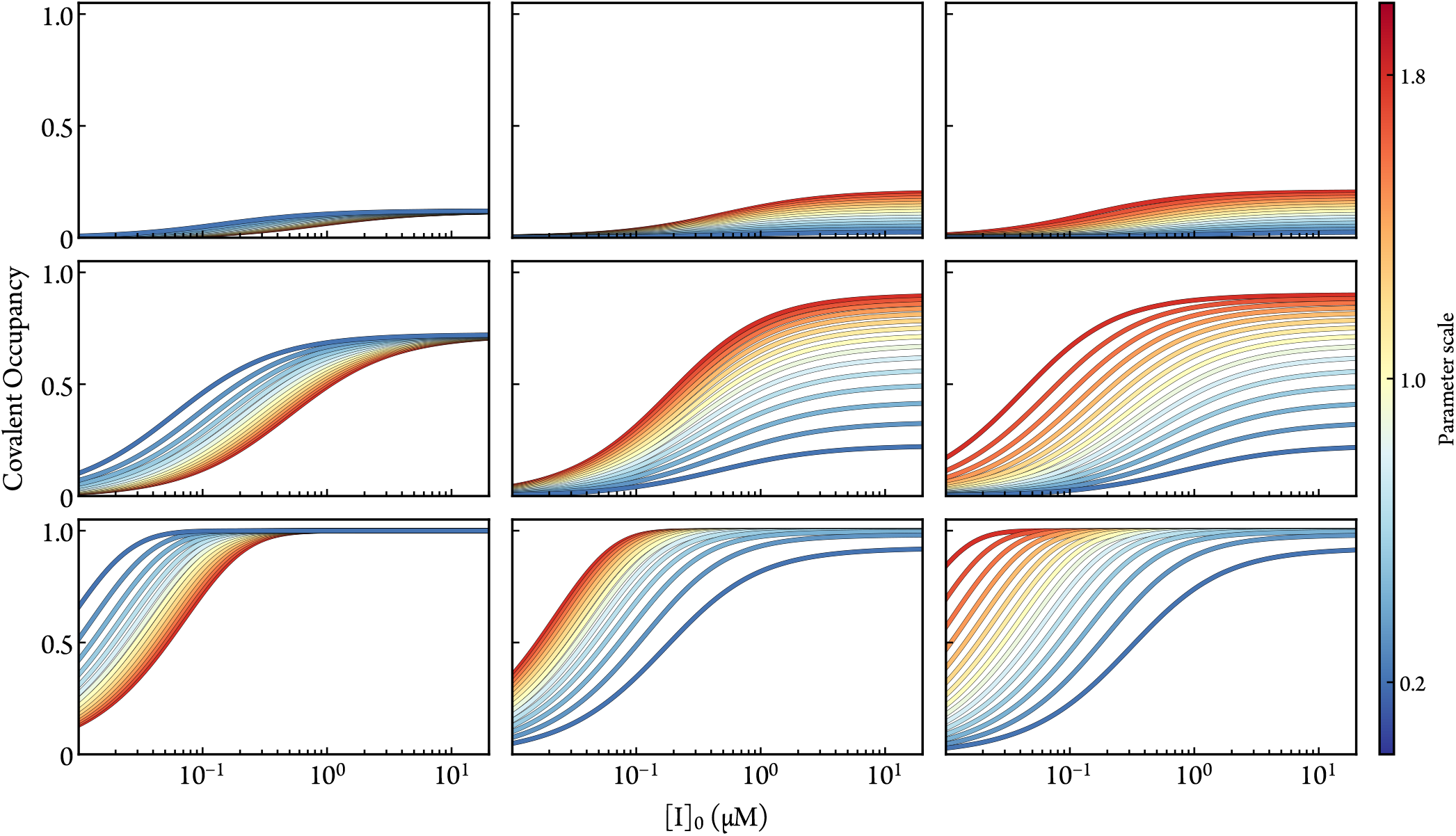
The variation of dose-response data (simulated by numerically solving the ODEs of Scheme 3) as a function of *k*_off_ (left column), *k*_inact_ (middle column), and *p*_r_ (right column) when *t*_DR_ is 0.7 (top row), 7.1 (middle row), and 71.1 (bottom row) hours. The default parameters are [E]_0_ = 1 nM, *k*_on_ = 1 × 10^−2^ µM^−1^s^−1^, *k*_off_ = 1 × 10^−2^ s^−1^, *k*_inact_ = 1 × 10^−4^ s^−1^, and *p*_r_ = 0.5.

The amplitude and inflection point of eq 16 taken together reveal the unique roles of *K*_d_, *p*_r_, and *k*_inact_. As shown in Figure 5, proportional variations of these simulation parameters distinctly affect the dose-response curves. The lack of complete covariance for these parameters suggests that fitting dose-dependent data with parameterized simulations is a viable strategy for empirical approximation of *K*_d_, *p*_r_, and *k*_inact_. If a set of dose-response curves (for different, *e*.*g*., experimental conditions or inhibitor modifications) is used in the fitting process, then one may discern the conditional impact on each parameter. If *p*_r_ is predicted with molecular simulations and held fixed, then *K*_d_ and *k*_inact_ could be approximated from a single dose-response curve.

The dose-response simulations also demonstrate the importance of a well-chosen timepoint. If *t*_DR_ is too short, as is shown in Figure 5 (top row), then the analysis of covalent occupancy is more obscured by absolute sources of error. If *t*_DR_ is too long and the reaction reaches 100% covalent occupancy, as is shown in Figure 5 (bottom row), then 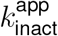 and *k*_*off*_ cannot be resolved in the fitting process.

By separating the contribution of rapid intermediates (*e*.*g*. conformations of the warhead or target residue, translation of the scaffold, protonation states) from the dissociation and inactivation steps, the kinetic comparisons between drug candidates become more precise, which should make it more straightforward to associate molecular characteristics with more potent and selective inhibition. This is especially beneficial when observed rates from reactivity or non-specific binding assays do not correlate with observed rates of inactivation or when the covalent action displays nonlinear complexities (as is the case for lysine-binders).^52,79^

While eq 16 is a useful approximation expressing the relationships between covalent occupancy and kinetic parameters, *in vitro* kinetics are more accurately represented with ODE-based simulations. This avoids the vague nature of the Hill coefficient *n* for fitting irreversible kinetics since noncooperative causes of a fitted *n* > 1, *e*.*g*. ultrasensitivity and significant depletion of [I], are unlikely to symmetrically affect the dose-response around the midpoint in the manner *n* is designed to modulate.^80,81^ ODE-based simulations also sidestep the assumptions and limitations of a pseudo-first order model such as the exponential growth factor,

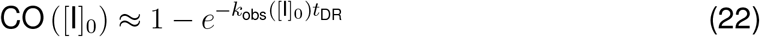

where *k*_obs_([I]_0_) uses eq 10. This extends to the dose-rate analysis using a time-dependent growth factor (eq 9) and eq 10.

If one globally fits parameterized simulations to multiple time-responses across varying doses—data required for dose-rate analysis anyway—then the fitting quality is no longer dependent on a single, well-chosen timepoint. The dose-dependence is included without the loss of information that occurs when an entire time course is compressed into a single value of *k*_obs_. Since ODE-based simulations accommodate low doses without concern for the validity of a first-order approximation, this strategy may enable a more accurate and practical (especially if *k*_obs_ would only plateau at unreasonably high doses) estimation of 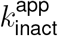 and 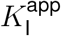 compared to the dose-rate workflow. The estimation of inactivation efficiency is also likely to be more accurate, because the initial slope of *k*_obs_([I]_0_) is susceptible to the nonlinearity of the bimolecular association step that emerges at low doses.

In ending this discussion, it is important to recall that the present analysis is meant to serve mainly as a conceptual illustration of how microscopic factors, that are not immediately related to the warhead’s chemistry, can impact the apparent observables such as the inhibition rate and residence time. Clearly, even with the expanded associated-unlinked intermediate state, Scheme 2 remains fairly simple and is unable to account for all the complex microscopic aspects of covalent inhibition encountered with different warheads. Expanding this type of analysis to realistic situations requires detailed atomic studies of the inhibition mechanism for specific cases. As an example, we previously represented the overall time-course of inhibition of BTK by ibrutinib, which targets a conserved cysteine residue, using an elaborate multi-state kinetic scheme based on the wealth of information extracted from QM/MM simulations.^52^ The mechanisms and reaction pathways of other cysteine-targeting covalent inhibitors could be examined using a similar strategy.^82,83^ Beyond cysteines, an understanding of covalent inhibition in the case of lysine-targeting warheads^84^ or other chemistries^22,23^ will require detailed atomic studies.

## Conclusion

A critical issue with the minimal two-step kinetic model, which represents the process of covalent inhibition as an association followed by formation of a chemical bond, is that it may obscure important microscopic factors that affect the apparent efficiency of a lead compounds. The propensity of the warhead and the target residue to adopt a reaction-ready configuration affects the apparent rate of inhibition but is not, in fact, specifically related to the warhead itself. The lack of clarity may on this issue and misinterpretation of experimental kinetic data can, in turn, misdirect the design strategy and pull a project away from a productive direction. To make judicious design decisions, it is critical to draw accurate lessons from experimental data. To this end, we explored a more realistic kinetic scheme, which can be used as a framework to incorporate information about the fluctuations of the associated-unlinked inhibitor generated from classical MD simulations.

Analysis based on numerical simulations showed that rapidly-equilibrating intermediate taking place on a short timescale can dramatically affect the apparent inhibition rate via the relative population of a reactive v.s. nonreactive configurations. Explicitly including the probability distribution of these states also refines the values of dissociation and inactivation rate constants, informing for example the amount of inactivation efficiency that can be attributed to only the warhead. Optimizing drug candidates with knowledge of the factors associated to the rapid fluctuations of the associated-unlinked inhibitor may offer a promising route for improving the apparent inactivation rate, while ignoring those could lead one to erroneously attribute a poor apparent inactivation efficiency to the warhead chemistry itself. Refining the evaluation with computational prediction of noncovalent behavior can provide a more precise evaluation of a drug’s potential and streamline drug optimization. Further work to develop kinetic frameworks incorporating microscopic states gleaned from classical MD simulations may be able to provide a more precise understanding of the molecular characteristics underlying apparent kinetic parameters.

## Acknowledgement

The authors thank Dr. Trayder Thomas and Prof. Raymond Moellering for the helpful discussion. This research was supported by the National Institutes of Health (NIH) via grant CA093577.

## Data and Software Availability

A Jupyter notebook for the kinetic modeling is available at https://github.com/RouxLab/covalent-kinetics-with-rapid-intermediates-2025. All data is deterministically generated therein. The in-house Python package for simulating the kinetics is available at https://github.com/kghaby/GeKiM.

